# The discovery potential of RNA processing profiles

**DOI:** 10.1101/049809

**Authors:** Amadís Pagès, Ivan Dotu, Joan Pallarès-Albanell, Eulàlia Martí, Roderic Guigó, Eduardo Eyras

**Affiliations:** Pompeu Fabra University (UPF), E08003 Barcelona, Spain.; Centre for Genomic Regulation (CRG), The Barcelona Institute of Science and Technology, E08003 Barcelona, Spain.; IMIM - Hospital del Mar Medical Research Institute. E08003 Barcelona, Spain.; Catalan Institution for Research and Advanced Studies (ICREA). E08010 Barcelona, Spain

## Abstract

Small non-coding RNAs are highly abundant molecules that regulate essential cellular processes and are classified according to sequence and structure. Here we argue that read profiles from size-selected RNA sequencing capture the post-transcriptional processing specific to each RNA family, thereby providing functional information independently of sequence and structure. We developed SeRPeNT, the first unsupervised computational method that exploits reproducibility across replicates and uses dynamic time-warping and density-based clustering algorithms to identify, characterize and compare small non-coding RNAs (sncRNAs) by harnessing the power of read profiles. We applied SeRPeNT to: a) generate an extended human annotation with 671 new sncRNAs from known classes and 131 from new potential classes, b) show pervasive differential processing between cell compartments and c) predict new molecules with miRNA-like behaviour from snoRNA, tRNA and long non-coding RNA precursors, potentially dependent on the miRNA biogenesis pathway. Furthermore, we validated experimentally four predicted novel non-coding RNAs: a miRNA, a snoRNA-derived miRNA, a processed tRNA and a new uncharacterized sncRNA. SeRPeNT facilitates fast and accurate discovery and characterization of small non-coding RNAs at unprecedented scale. SeRPeNT code is available under the MIT license at https://github.com/comprna/SeRPeNT.

## Background

Small non-coding RNAs (sncRNAs) are highly abundant functional transcription products that regulate essential cellular processes, from splicing or protein synthesis to the catalysis of post-transcriptional modifications or gene expression regulation (1). Major classes include micro-RNAs (miRNAs), small nucleolar RNAs (snoRNAs), small nuclear RNAs (snRNAs) and transfer RNAs (tRNAs). Developments in high-throughput approaches have facilitated their characterization in terms of sequence and structure (2–4) and have led to the discovery of new molecules in diverse physiological and pathological contexts. However, the function of many of them remains unknown (5, 6); hence their characterization is essential to understand multiple cellular processes in health and disease.

Sequence and structure are traditionally used to identify and characterize small non-coding RNAs (7, 8). Although sequence is a direct product of the sequencing technology, structure determination is still of limited accuracy and requires specialized protocols (3, 4, 9). On the other hand, extensive processing is a general characteristic of non-coding RNAs (10–12). The best-characterized cases are miRNAs, which are processed from precursors and preferentially express one arm over the other depending on the cellular conditions (13, 14). Furthermore, snoRNAs and tRNAs can be processed into smaller RNAs, whose function is often independent of their precursor (10, 15–18). These findings suggest that a new path to systematically characterize RNA molecules emerges through the genome-wide analysis of their sequencing read profiles.

Here we argue that sequencing profiles can be used to directly characterize the function of small non-coding RNAs, in the same way that sequence and structure have been used in the past. We report here on SeRPeNT, a fast and memory efficient software for the discovery and characterization of known and novel classes of small non-coding RNAs exploiting their processing pattern from small RNA sequencing experiments. SeRPeNT is the first unsupervised algorithm to classify and characterize sncRNAs. As opposed to previous supervised methods that necessarily rely on known annotations, SeRPeNT is capable of grouping sncRNAs into families without the need of previous annotation, and therefore has the potential to discover new classes of sncRNAs. We applied SeRPeNT to generate an extended human annotation with 671 new RNAs from known classes and 131 from new potential classes. We further showed these sncRNAs to have pervasive differential processing between cell compartments and predict new miRNA-like molecules that are potentially processed from different RNA precursors, including snoRNAs, tRNAs and long non-coding RNAs. Finally, we validated experimentally four novel non-coding RNAs predicted by SeRPeNT, highlighting the power of SeRPeNT for the discovery and characterization of small non-coding.

## Methods

Using multiple size-selected small (<200nt) RNA sequencing (sncRNA-seq) experiments mapped to a genome reference, SeRPeNT enables the discovery and characterization of known and novel small non-coding RNAs (sncRNAs) through three operations: *profiler*, *annotator* and *diffproc*, which can be used independently or together in a pipeline (Fig. 1). Initially, sncRNA read profiles are calculated from the mapped sncRNA-seq reads, and filtered according to the reproducibility between replicates, and to the length and expression constraints given as input (Fig. 1A). Pairwise distances between profiles are calculated as a normalized cross-correlation of their alignment calculated using a time-warping algorithm (Fig. 1B). Profiles are clustered into families according to pairwise distances using an improved density-based clustering algorithm (Fig. 1B). Novel profiles are annotated using the class label from known profiles in the same cluster if available by majority voting (Fig. 1C). Additionally, SeRPeNT allows the identification of differential processing of sncRNAs between two conditions, independently of their expression change (Fig. 1D).

**Figure 1.**
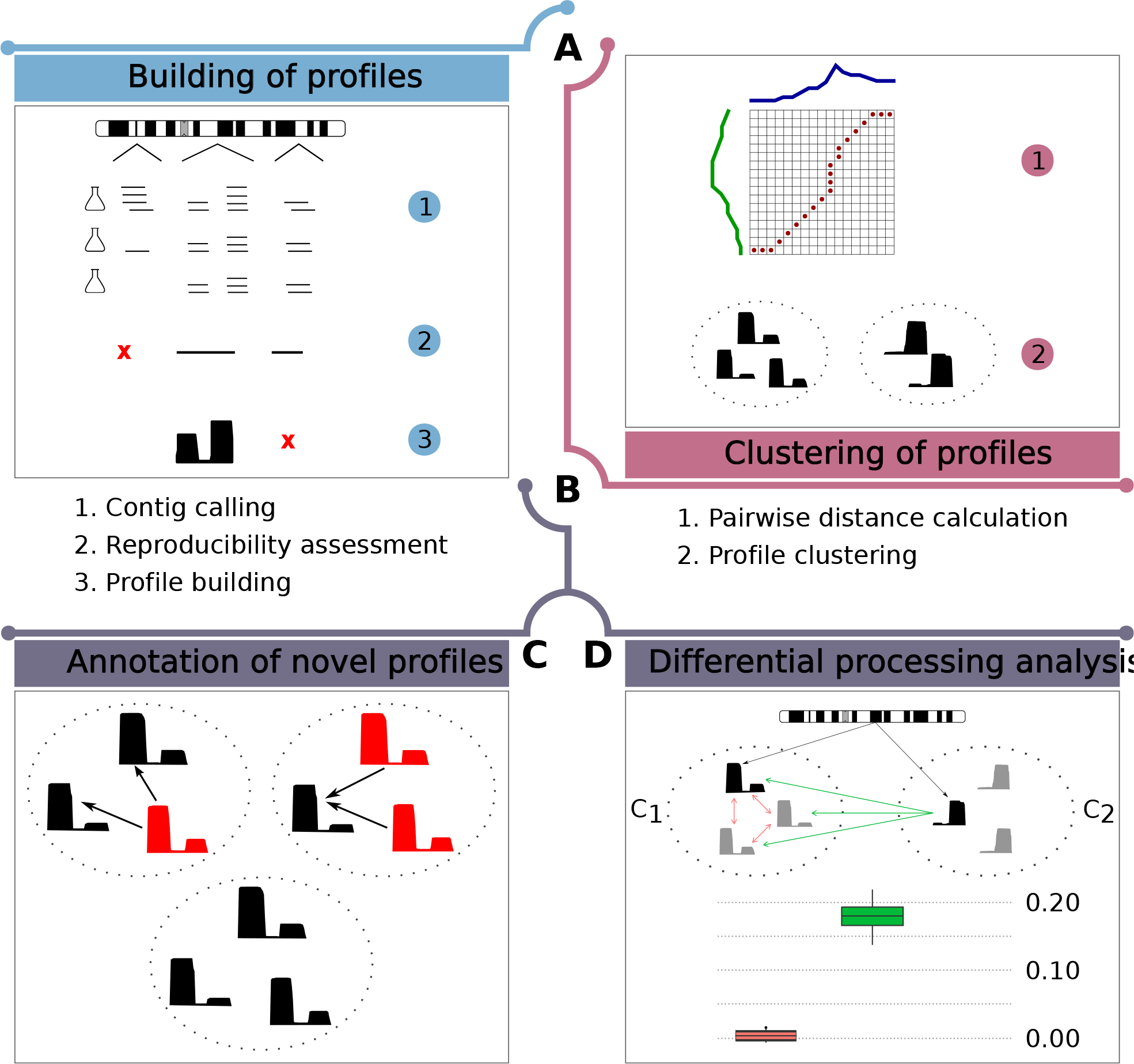
Overview of SeRPeNT. Overview of the operations performed by the SeRPeNT: **(A)** Building of profiles from short RNA-Seq reads mapped to the genome using reproducibility across replicates. A profile is a collection of reads overlapping over a given genomic locus and can be regarded as a vector where each component contains the number of reads at each nucleotide. (**B**) Density-based clustering of profiles based on pairwise distances calculated with a dynamic time-warping algorithm. **(C)** Annotation of novel profiles using majority vote in clusters. **(D)** Differential processing calculation. The distribution of distances between a profile and its cluster sisters in one condition cluster (C_1_) and across conditions (C_2_) are compared (panel below). Differential processing is determined in terms of a Mann-Whitney U test and a fold-enrichment (Supplementary Fig. 2).

### Profile building from aligned short RNA-Seq reads

The tool *profiler* uses as input one or more small RNA sequencing replicates in BAM format. Consensus read contigs are built by pooling reads that overlap on a genomic region and that are at a distance smaller than a user-defined threshold. Each contig is scored per individual replicate by counting the number of reads mapped within its boundaries and reproducibility is measured across all the biological replicates using either a non-parametric irreproducibility detection rate (NP-IDR) (19) or the simple error ratio estimate (SERE) (20). NP-IDR determines the reproducibility of a contig in one or more replicates with similar sequencing depths, whereas SERE compares the observed variation in the raw number of reads of a contig to an expected value, accounting for the variation in read depth across replicates. For all analyses of reproducibility in this work we used NP-IDR with cut-off of 0.01. Contigs that do not pass the user-defined cutoff of reproducibility are discarded from further analysis. For each of the remaining contigs, a profile is built by counting the number of reads per nucleotide in the genomic region delimited by the contig boundaries (Fig. 1A). SeRPeNT defines each sncRNA as a genomic region and a vector of raw read counts, or heights, of length equal to the number of nucleotides spanned by this genomic region. Profiles are additionally trimmed at the 3’-end positions when heights are either below 5 reads or below 10% of the highest position, but not when having more than 20 reads. Only profiles of lengths between 50 and 200 nucleotides, and of minimum height 100 in pooled replicates, were considered. All these parameters can be configured on SeRPeNT command line. The consistency of sncRNA profiles across multiple experiments is determined by calculating a normalized entropy of the different labels for the same sncRNA locus across experiments (Supplementary Materials and Methods).

### sncRNA profile clustering

The *annotator* tool assigns a distance between each possible pair of profiles resulting from the previous step. This distance is computed with a novel algorithm (described in Supplementary Figure S1) based on dynamic time-warping (21, 22) (Fig. 1B). This algorithm finds the optimal alignment between two profiles by placing the heights of a pair of profiles along the axes of a grid, representing alignments as paths through the grid cells, and finding the path with maximum normalized cross-correlation score across them. Given a pair of profiles of the same length *A = (a_1_, …, a_n_)* and *B = (b_1_, …, b_n_)*, where *a_i_* and *b_i_* are the heights of nucleotide *i* in profile *A* and *B*, respectively, the cross-correlation score between *A* and *B* is defined as:

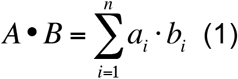

and the normalized cross-correlation score as:

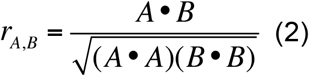

The optimal alignment maximizes the normalized cross-correlation score between the two profiles. Given two profiles *S = (s_1_, …, s_n_)* and *Q = (q_1_, …, q_m_)* of length *n* and *m* nucleotides respectively, each position *(i, j)* in the dynamic programming matrix *D* stores a vector of three values *D*(*i,j*) = (*x, y, z*) such that they maximize the value 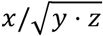 in formula (2) amongst all the possible partial alignments between *S_i_* and *Q*_*j*_, where *S*_*i*_ = (*s*_1_, …, *s_i_*) and Q_*j*_ = (*q*_1_, …, *q*_j_) are the profiles spanning the first *i* and *j* nucleotides of the profiles *S* and *Q*. The dynamic programming equation is then defined as:

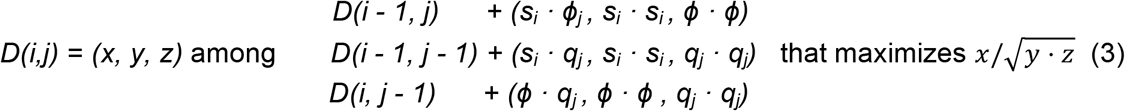

where ϕ represents a negative Gaussian white noise function used to penalize an expansion or contraction in the alignment. When applied to a profile *S*, *ϕ(S)* returns a random negative value taken from a uniform distribution with mean and standard deviation defined by *S*.

Once all the pairwise distances are calculated, profiles are clustered using a modified version of a density-based clustering algorithm (23) (described in Supplementary Figure S2A). The clustering algorithm is based on the assumption that clusters are formed by points surrounded by a high density of data points of lower local density and lie at large distance from other profiles of high local density. For each profile *i* we defined its local density *ρ_i_* as follows:

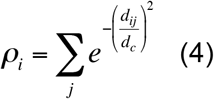

where *d_ij_* is the distance between profiles *i* and *j*, and *d_c_* is an optimal distance that determines the size of the neighborhood of a profile. The optimal distance *d_c_* is calculated using a data field calculated from all profiles (24, 25) (algorithm described in Supplementary Figure S2B). Once the optimal *d_c_* is obtained, the profile with the highest local density is identified, and this profile and all the profiles that are within distance *d_c_* are assigned to a cluster. We introduced a novel step in the clustering in which all the profiles that have already been clustered are removed before the next iteration step. In the next step, a new *d_c_* value is then calculated with the remaining clusters and new local densities are calculated to identify the cluster with highest density, and so on. The algorithm stops when only singletons are produced or when the calculated optimal value for *d_c_* is higher than 0.02. This value represents the maximum distance we allow to start building a cluster from a profile with the highest local density.

### Profile annotation

The *annotator* tool performs the sncRNA profile annotation. Every detected profile that overlaps an annotated short non-coding RNA is marked as known and labeled with the corresponding class label (e.g. H/ACA snoRNA). The minimum overlap amount required between the sncRNA profile and the annotated RNA can be defined by the user. Profiles that do not overlap with any annotation or do not satisfy the overlapping requirements are marked as unknown. For each cluster with two or more profiles, the different labels from all the known profiles are counted, and all the unknown profiles within the cluster are labeled by majority vote with the most abundant label (Fig. 1C). In case of a tie, the label of the closest profile is assigned. All the remaining profiles are denoted as unlabeled. Clustered unlabeled profiles represent a coherent group of multiple profiles, and hence potentially indicate a novel sncRNA class.

### Differential processing analysis

Differential processing is calculated for each sncRNA from the pairwise distance distributions with sister sncRNAs from the same cluster in either condition. Profiles are considered as differentially processed according to the fold-change and significance of the change. The *diffproc* tool assesses if a profile *P_a_* in a particular condition *C_1_* shows a different processing pattern *P_b_* in another condition *C_2_* (Fig. 1D). A pair of profiles *P_a_* and *P_b_* from conditions *C_1_* and *C_2_*, respectively, such that their reference coordinates overlap as described above, are compared as follows. Given *K_a_* the cluster in condition *C_1_* that contains the profile *P_a_* and *K_b_* the cluster in condition *C_2_* that contains the profile *P_b_*, *diffproc* calculates all the pairwise distances *D_ab_* between *P_a_* and all the profiles in *K_b_*, and the pairwise distances *D_b_* between profiles in *K_b_* (Fig. 1). These two distance distributions are then compared using a one-sided Mann-Whitney *U* test and a fold-change is calculated as the ratio of the medians between both distributions. The same method is applied to profile *P_b_* and cluster *K_a_*. *P_a_* and *P_b_* are then reported as differentially processed if both tests are significant according to the p-value and fold-change cutoffs defined by the user. When there are not enough cases to perform a Mann-Whitney U test, only the fold-change is taken into account.

### Accuracy analysis and experimental validation

Details about the accuracy analysis and the experimental validations are available in the Supplementary Material.

### Software

SeRPeNT is written in C. The source code is available at https://github.com/comprna/SeRPeNT Code and makefiles to reproduce the analyses described in this manuscript are available at https://github.com/comprna/SeRPeNT-analysis

## Results

### Fast and accurate discovery of small non-coding RNAs

We assessed the accuracy of SeRPeNT by performing a comparison against BlockClust (26), an unsupervised method that also predicts known small non-coding RNA families from small RNA sequencing (sncRNA-seq) data. We evaluated the accuracy to detect known miRNAs, tRNAs, and snoRNAs from the Gencode annotation (27) using the same procedure and dataset used by Videm et al. (26) (Supplementary Materials and Methods). SeRPeNT shows overall similar precision for miRNAs (0.858) and tRNAs (0.855), and a dramatic improvement of the precision for snoRNAs (0.922) (Supplementary Table S1A). Of note, although BlockClust was benchmarked by Videm et al. using only C/D-box snoRNAs only, we benchmarked SeRPeNT using also H/ACA-box snoRNAs. Notably, SeRPeNT analysis took ~3 minutes and less than 200Mb of RAM in a single core AMD Opteron 64 with 4Gb of memory. In contrast, the same analysis with BlockClust, which included the execution of Blockbuster (28), took ~15 minutes and used nearly 30Gb of memory. Additionally, we compared the performance of SeRPeNT against the supervised version of BlockClust and against DARIO (27), using a cross-fold validation approach (Supplementary Figure S3). Using small RNA-Seq data from MCF-7 cells (8) (GSM769510) for the three methods, SeRPeNT shows overall higher precision in all tested sncRNA families (Supplementary Table S1B). Importantly, as opposed to the supervised methods, SeRPeNT did not use the annotation to group sncRNAs profiles.

We also assessed the accuracy of SeRPeNT differential processing operation *diffproc* by analyzing the differential expression of miRNA arms and arm-switching events in miRNAs between normal and tumor liver tissues (29). From the 49 miRNAs tested, 41 passed our filters of reproducibility and clustered with other sncRNAs. Imposing a significance threshold of p-value < 0.01 and a fold-change of at least 2.5 (Supplementary Figure S5), SeRPeNT identified as differentially processed 10 out of 24 miRNAs described to exhibit different 5’-arm to 3’-arm expression ratio (29), including 4 out of 5 arm-switching events (Supplementary Figure S6). Moreover, only 1 out of the remaining 17 miRNAs that did not exhibit a difference in 5’-arm to 3’-arm expression ratio was identified as differentially processed by SeRPeNT. We further compared SeRPeNT against RPA (30), a recent method for differential processing analysis, using sncRNA-seq data from 9 cell lines (31). SeRPeNT detected many more differentially processed events, with a moderate overlap with RPA predictions (Supplementary Figure S7). Notably, for this analysis SeRPeNT took 2 hours in a single core AMD Opteron 64 with 4Gb of memory, whereas RPA took about 10 hours in a cluster of 32 cores each having 8 Gb of RAM.

### An extended annotation of small non-coding RNAs in human

We decided to exploit SeRPeNT to produce an extended annotation of small non-coding RNAs in human. We applied SeRPeNT *profiler and annotator* tools to sncRNA-seq data from 9 cell lines (31) (Supplementary Table S2). We observed a higher proportion of known RNAs compared to novel sncRNAs, with an increase of novel sncRNAs in samples sequenced at a higher depth: A549, IMR90, MCF-7 and SK-N-SH (Fig. 2A). We further measured the accuracy of SeRPeNT in recovering known sncRNA classes using cross-fold validation in these datasets and found an overall high accuracy consistently across all cell lines (Supplementary Table S3), except for snRNAs, probably due to their broad differences in structural features and processing patterns (12). Additionally, in the cross-fold validation SeRPeNT did not annotate on average about 30% of all the profiles detected in known scnRNAs from Gencode, as they either were in clusters with only unlabelled profiles or because they were singletons. Importantly, the accuracy values were robust when running SeRPeNT with different parameters for minimum expression, reproducibility value between replicates, minimum length of sncRNA profiles or spacing between profiles, and using different sequencing depths (Supplementary Table S4).

**Figure 2.**
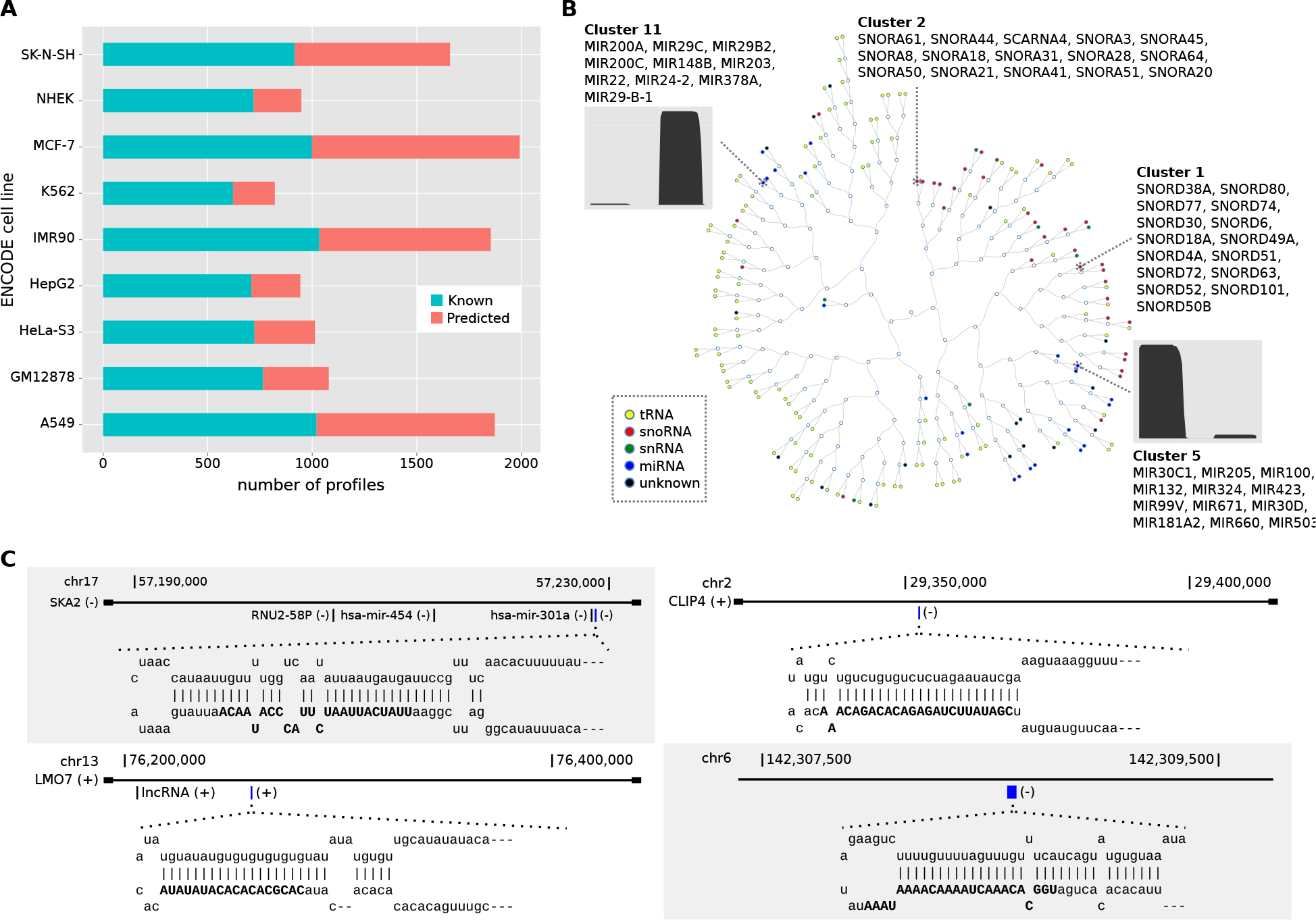
Extended annotation derived from ENCODE cell lines. **(A)** Number of known and novel sncRNAs across 9 ENCODE cell line dataset. **(B)** Hierarchical clustering representation of the clusters obtained for the NHEK cell line. Distance between clusters is calculated by averaging all the distances between profiles from both clusters. Colored circles represent clusters of sncRNAs at the leaves of the tree labeled by class. Empty circles represent internal nodes of the tree. The read profiles in clusters 5 and 11 are for one of its members, for which we plot the number of reads per nucleotide in the sncRNA. **(C)** Genomic loci and graphical representation of the hairpins for four predicted novel microRNAs. The predicted mature microRNAs are highlighted in blue in the corresponding gene loci: miRNA chr17:57228820-57228919:- (upper left) at the *SKA2* locus, miRNA chr2:29352292 -29352349:- (upper right) at the CLIP4 locus, miRNA chr13:76258915-76258974:+ (lower left) at the *LMO7* locus, and miRNA chr6:142308575-142308638:- (lower right) at an intergenic region.

We annotated new sncRNAs with SeRPeNT and obtained a total of 4,673 non-unique sncRNAs across all tested cell lines that were not in the Gencode annotation (Supplementary Table S5). We were able to assign a label to 2,140 of them. From the remaining 2,533 unlabeled sncRNAs, 323 formed 92 clusters with three or more unlabeled profiles per cluster, suggesting possible new classes of non-coding RNAs with a coherent processing pattern. We called these *clustered uncharacterized* RNAs (cuRNAs) and kept them for further study. Interestingly, some known and predicted sncRNAs with the same class labels were grouped into different clusters, indicating subfamilies. For instance, SeRPeNT separated C/D-box and H/ACA-box snoRNAs according to their processing profiles (clusters 1 and 2 in Fig. 2B), and separated miRNAs into subtypes according to their different arm-processing patterns (clusters 5 and 11 in Fig. 2B). Thus SeRPeNT identifies functional families and subfamilies of non-coding RNAs in a scalable and robust way, independently of the granularity of the available annotation.

We established the consistency of the sncRNAs across the multiple experiments using an entropy measure of the label assignment across cell lines (Supplementary Materials and Methods), producing a total of 929 unique novel sncRNAs, 787 from the major classes (79 miRNAs, 475 snoRNAs, 82 snRNAs and 151 tRNAs) plus 142 cuRNAs, the majority of them being expressed in only one cell line (Supplementary Figure S7). These, together with the sncRNAs annotated in Gencode, conformed an extended catalogue of small non-coding RNAs in the human genome reference (Supplementary Table S6), also available in GTF format as Supplementary file.

From the 79 newly predicted miRNAs, 37 (46.8%) were confirmed as potential miRNA precursors using FOMmiR (32) (Supplementary Table S7). Moreover, 39 (49.3%) of these novel miRNAs overlapped with AGO2-loaded small RNAs from HEK293 cells (33). In contrast, from 3109 known miRNAs in the Gencode v19 annotation, 951 (30,59%) overlapped with AGO2-loaded small RNAs (Fisher’s exact test p-value = 1.14e-3, odds-ratio = 2.01) (Supplementary Tables S6 and S7). To further characterize these miRNAs, we searched for sequence and secondary structure similarities in Rfam using Infernal (34, 35), with threshold e-value < 0.01 (Supplementary Materials and Methods). We found that 23 of them had a hit to a known miRNA family (Supplementary Table S7). Repeating these analyses for the other new sncRNAs we found 47 snoRNA and 15 tRNAs with a hit to an Rfam family, from which 3 snoRNAs and 4 tRNAs had a hit to a family of the same class predicted by SeRPeNT (Supplementary Table S6). The rest of predicted sncRNAs did not have any hit to Rfam. We further compared the predicted sncRNAs from our extended annotation with DASHR (6), the most recently published database of human small non-coding RNAs, and with a compendium of human miRNAs from a recent study using multiple samples (36). We found that 802 out of the 929 predicted sncRNAs (51 miRNAs, 430 snoRNAs, 69 snRNAs, 121 tRNAs and 131 cuRNAs) were not present in those catalogues. In particular, four of the newly predicted miRNAs that had a hit to an Rfam miRNA family and were confirmed as potential miRNA precursors with FOMmiR were not present in these previous catalogues (6, 36) (Fig. 2C). We further checked the overlap of cuRNAs with CAGE data from The FANTOM5 project (37) (Methods). From the 142 cuRNAs in the extended annotation, 32 of them overlapped with CAGE profiles in the same strand. Moreover, for 27 of these 32 (84.3%) the 5’ end of the cuRNA overlaps with the CAGE profile (Supplementary Table S8).

### SeRPeNT uncovers new RNAs with potential miRNA-like function

SeRPeNT analysis on individual cell lines identified a cluster that grouped together snoRNA SCARNA15 (ACA45) with 2 miRNAs in NHEK, and a cluster that grouped snoRNA SCARNA3 with several miRNAs and a tRNA in A549 (Supplementary Table S9) in agreement with a previous study showing that these snoRNAs can function as miRNAs (15). The clusters obtained with SeRPeNT in cell lines provided additional evidence of 6 other snoRNAs that grouped with miRNAs: SNORD116, SNORA57, SNORD14C, SNORD26, SNORD60 and SNORA3 (Supplementary Table S9), suggesting new snoRNAs with miRNA-like function. Interestingly, we also found 7 clusters with a majority of miRNAs that included annotated tRNAs: tRNA-Ile-GAT, tRNA-Glu-GAA, tRNA-Gly-CCC, tRNA-Ala-AGC and tRNA-Leu-AAG. In particular, tRNA-Ile-GAT-1-1 clusters with miRNAs in 3 different cell lines: MCF-7, A549 and SK-N-SH, suggesting new tRNAs with miRNA-like function (10, 38). These results support the notion that sncRNA read-profiles facilitate the direct identification of functional similarities without the need to analyze sequence or structure.

To search for new cases of miRNA-like non-coding RNAs in the extended annotation, we tested their potential association with components of the canonical miRNA biogenesis pathway, using sncRNA-seq data from controls and individual knockouts of *DICER1*, *DROSHA* and *XPO5* (39) (Supplementary Materials and Methods). We validated the dependence of a number of known and predicted miRNAs on these three factors (Fig. 3A) (Supplementary Figures S9 and S10) and recovered the previously described dependence of ACA45 and SCARNA3 with *DICER1* (15). Additionally, we found 18 sncRNAs predicted as snoRNAs with similar behaviour upon *DICER1* knockout (Fig. 3B). Interestingly, 14 out of 20 *DICER1*-dependent snoRNAs did not show dependence on *DROSHA*, including ACA45 and SCARNA3, in agreement with previous findings (15, 39) (Supplementary Figure S8) (Supplementary Table S7). We also found a strong dependence on *DICER1* for 128 tRNAs, 82 of which changed expression in the direction opposite to most miRNAs, suggesting that they may be repressed by DICER (Fig. 3C). Further, 4 cuRNAs showed similar results to miRNAs, suggesting some association with the miRNA biogenesis machinery (Supplementary Figure S10) (Supplementary Table S7). Although they were not confirmed as potential miRNA precursors using FOMmiR, 2 of these miRNA-like cuRNAs overlapped with the protein-coding genes *SEC24C* and *DHFR* (Supplementary Figure S10).

**Figure 3.**
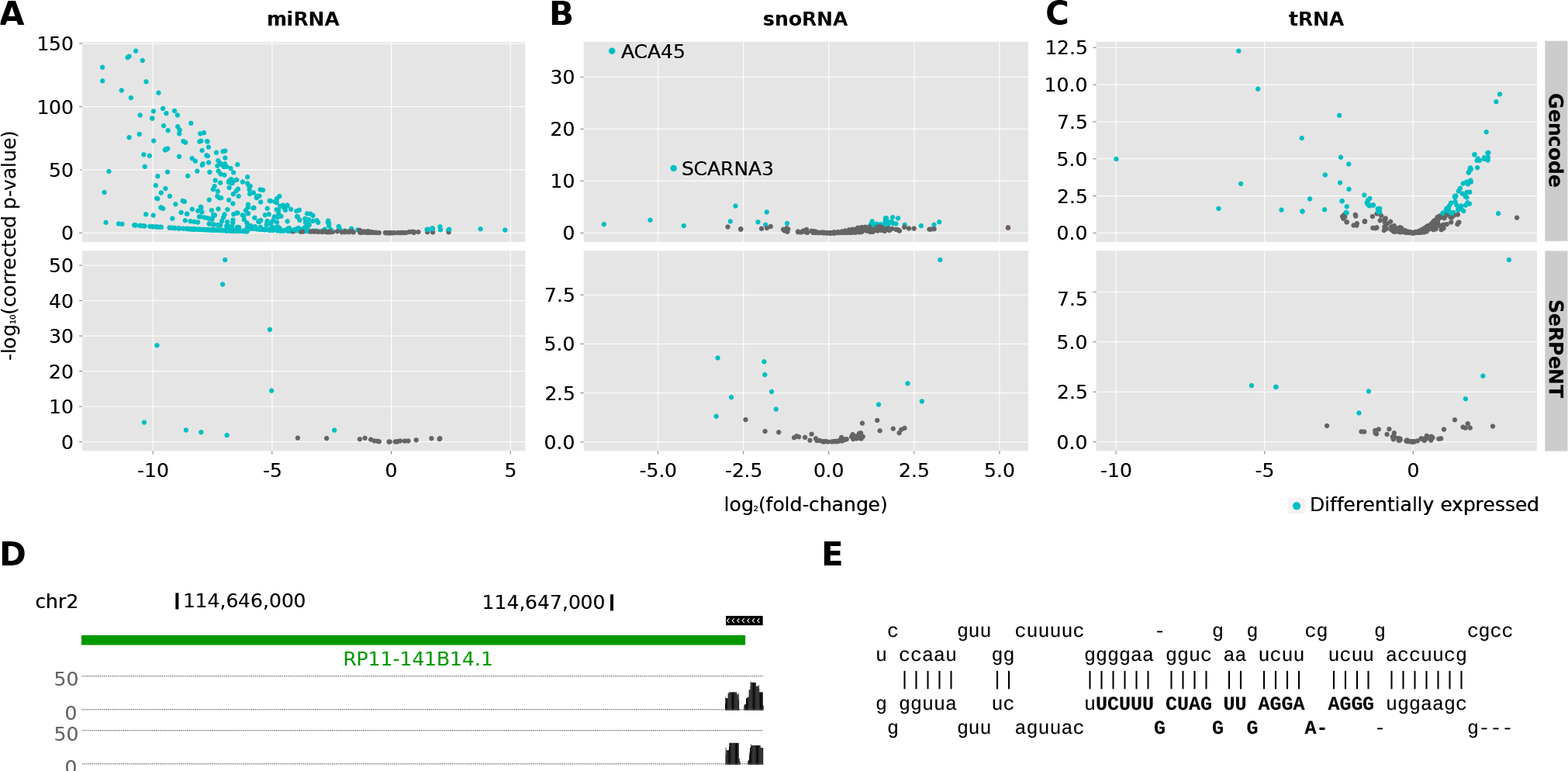
Detection of miRNA-like sncRNAs. Differentially expressed sncRNAs (blue) from extended annotation in the comparison between *DICER1* knockout and control experiments in human HCT116 cell lines for **(A)** miRNAs, **(B)** snoRNAs and **(C)** tRNAs. The analyses for the knockout of *DROSHA* and *XPO5* are available as Supplementary Figures. **(D)** Representation of a novel miRNA detected by SeRPeNT (depicted as a read profile) whose precursor is the lncRNA RP11-141B14.1 (depicted as a green line). Profiles for both replicates are included. **(E)** Secondary structure prediction of the miRNA precursor by FOMmiR.

Certain long non-coding RNAs (lncRNAs) are known to act as precursors of miRNAs (40, 41) and tRNAs (42). We thus analyzed whether the new sncRNAs could originate from lncRNAs. We found that 8 miRNAs, 16 snoRNAs, 7 tRNAs and 4 cuRNAs overlapped annotated lncRNAs (Supplementary Table S7). These lncRNAs included *MALAT1*, which we predicted to produc 2 miRNAs, 2 tRNAs and 1 cuRNA. Additionally, 3 of the miRNAs predicted and confirmed with FOMmiR were found on the lncRNAs *MIR100HG*, *CTD-23C24-1*, and *RP11-141B14.1*. From these, the new miRNA in *RP11-141B14.1* is not present in recent miRNA catalogues (Figs. 3D and 3E). As the processing from lncRNAs is a recognized biogenesis mechanism for certain small non-coding RNAs, these results provide further support for the relevance of the newly predicted sncRNAs in our extended annotation.

### Pervasive differential processing of non-coding RNAs between cell compartments

To further characterize the extended sncRNA annotation defined above, we studied their differential processing between four different cell compartments: chromatin, nucleoplasm, nucleolus and cytosol for the cell line K562 using replicated data (31) (Supplementary Table S2). The majority of sncRNAs from the extended annotation showed expression in one or more cell compartments: 599 in chromatin, 763 in cytosol, 554 in nucleolus and 651 in nucleoplasm. The majority of sncRNAs in cytosol are tRNAs (45%), followed by miRNAs (15%). Although tRNAs were enriched in the cytosol (Fisher’s one-sided test p-value < 0.001), they were abundant in all four cell compartments (Supplementary Table S10). This is compatible with tRNA biogenesis, which comprises early processing in the nucleolus and later processing in the nucleoplasm before export to the cytoplasm (43). In contrast, miRNA clusters appeared almost exclusively in the cytosol (Fisher’s one-sided test p-value < 0.001) and were coherently grouped into large clusters (Fig. 4A) (Supplementary Table S10). On the other hand, snoRNAs were enriched in the nucleolus (Fisher’s one-sided test p-value < 0.01), accounting for 38% of the found profiles. Interestingly, snoRNAs were also enriched in the chromatin compartment (Fisher’s one-sided test p-value <0.001) accounting for 23% of the sncRNAs found there, suggesting new candidates for their recognized role on establishing open chromatin domains (44). Finally, snRNAs and cuRNAs appeared at low frequency in most compartments (Supplementary Table S10). We applied SeRPeNT *diffproc* operation for each pair of compartments, using fold-change ≥ 2.5 and p-value < 0.01. A large proportion of snoRNAs showed differential processing from the nucleus and nucleolus, where they exert their function, to the rest of cellular compartments (Fig. 4B). On the other hand, only 4 of the cuRNAs identified showed expression in at least two compartments, nucleolus and cytosol, and 3 of them showed differential processing. Overall, tRNAs showed the largest proportion of differentially processed profiles between the cytosol and the different nuclear compartments (Fig. 4B and Supplementary Table S11). Many of these tRNAs showed a more prominent processing in the cytosol from the 30-35nt part of their 3’ part (Fig. 4C and Supplementary Figure S11), also called tRNA halves (45, 46).

**Figure 4.**
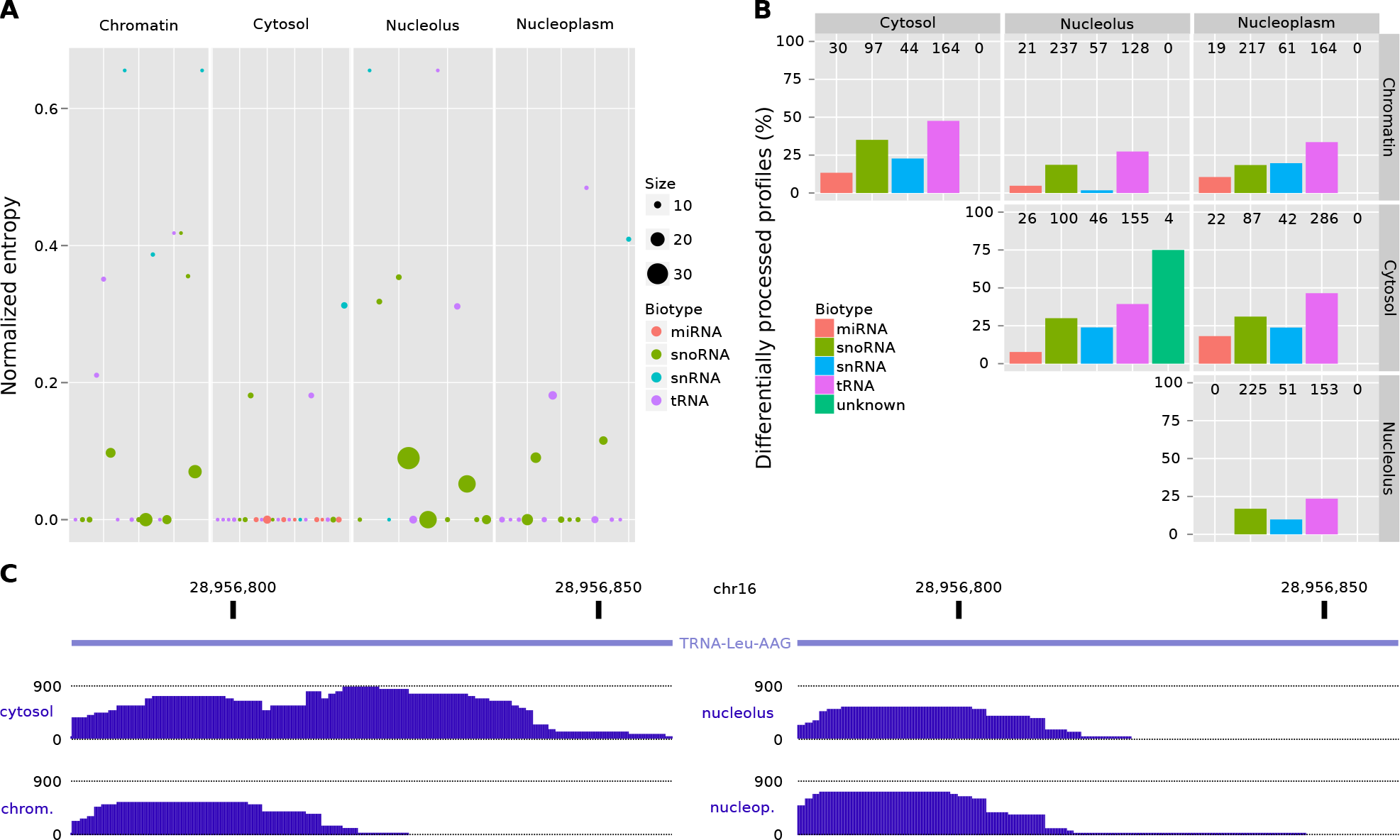
Differential processing across ENCODE cell compartments. **(A)** Representation of clusters containing 5 or more sncRNAs across all four ENCODE cell compartments. The size of the points represents the number of sncRNAs from the extended annotation contained in the cluster. The normalized entropy (y axis) represents the purity of a cluster (Supplementary Materials and Methods), the lower the entropy, the higher the purity of the cluster. **(B)** Proportion of profiles from the extended annotation that are differentially processed between cellular compartments separated by non-coding RNA family (y axis). Numbers at the top of the bars represent the total number of profiles detected in both compartments. **(C)** Representation of the read profiles for the tRNA-Leu-AAG transfer RNA showing abundant processing of the 3’-half in the cytosol compared to the chromatin compartment. The plot represents the number of reads per nucleotide in the same scale for each compartment.

### Experimental validation of novel short non-coding RNAs

To validate our findings, we decided to test experimentally four of the newly predicted sncRNAs that had no sequence similarity with any other genomic position using four different cell lines: SH-SY5Y, MCF7, MCF10 and HeLa (Fig. 5 and Supplementary Fig. S12) (Supplementary Tables S12 and S13) (Supplementary Methods). We selected a novel miRNA (chr17:57228820-57228919:-) that we predicted in the intron of *SKA2* (Fig. 2) (Supplementary Table S6). This candidate miRNA clustered with other miRNAs and was predicted to belong to the Rfam family RF00906. Using sequence specific primers we detected expression of this miRNA by qPCR in HeLa and SH-SY5Y cells (Fig. 5 and Supplementary Fig. S12). We detected this miRNA with SeRPeNT using small-RNA-seq from the same SH-SY5Y and MCF7 cells used for experimental validation (47) and using small-RNA-seq from ENCODE for HeLa (Fig. 5 and Supplementary Fig. S12). We also tested a novel miRNA that we predicted to be processed from the H/ACA snoRNA SNORA3 (chr16:2846409-2846473:-) (sno-miRNA in Fig. 5) (Supplementary Table S9). Using sequence specific primers, we detected expression of this miRNA by qPCR in HeLa and SH-SY5Y cell lines (Fig. 5). This miRNA was also detected with SeRPeNT using small RNA-seq data for the same SH-SY5Y cells and from ENCODE HeLa cells (Fig. 5 and Supplementary Fig. S12). Additionally, we tested experimentally a clustered-uncharacterized RNA (cuRNA) (chr10:75526203-75526253:+) that we had detected in SK-N-SH and IMR90 cells (Supplementary Table S6), and which we measured to be dependent of DROSHA, DICER and XPO5 (Supplementary Table S7). This cuRNA was predicted with SeRPeNT to be lowly expressed in HeLa and SH-SY5Y, but it was only detected by qPCR in HeLa (Figure 5 and Supplementary Figure S12). Finally, we tested a processed tRNA (p-tRNA) (chr1:145396847-145396952:-) that was predicted to be cytosol-specific in K562 cells and had differential processing with respect to the nucleolus and chromatin compartments (Supplementary Table S11). We validated this p-tRNA in all four cell lines used for validation. Moreover, we detected the p-tRNA with SeRPeNT using small RNA-seq from the same SH-SY5Y, MCF7 and MCF10 cells used for experimental testing (47) and from ENCODE HeLa cells (Figure 5 and Supplementary Figure S13).

**Figure 5.**
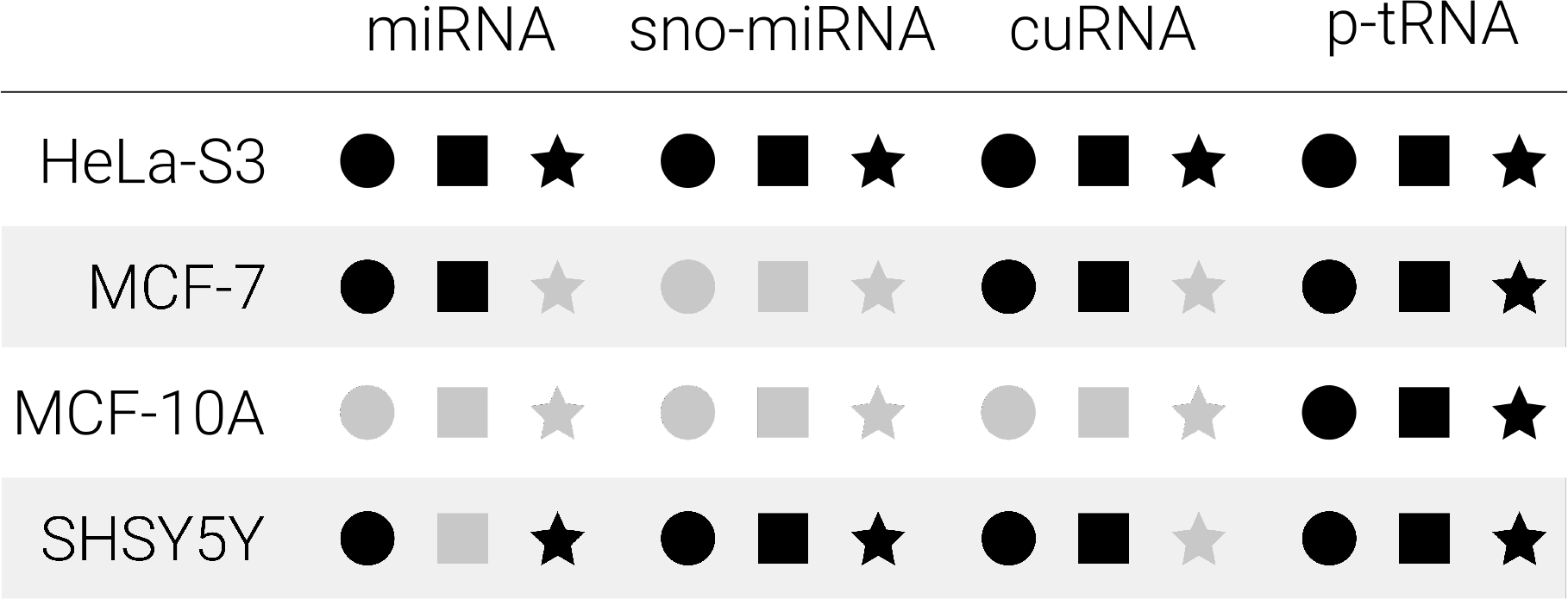
Experimental validation of novel sncRNAs. Experimental validation of four predicted sncRNAs in the cell lines HeLa, SH-SY5Y, MCF10A, MCF7, SH-SY5Y and HeLa. We tested a predicted miRNA (miRNA), a miRNA predicted to be derived from the H/ACA snoRNA SNORA3 (sno-miRNA), a clustered uncharacterized RNA (cuRNA) and a processed tRNA (p-tRNA). For each sncRNA and each cell line, we indicate whether it was detected by SeRPNeNT (black circle), whether its measured expression was RPM (reads per million) > 1 (black triangle), and whether it was validated by qPCR (black star), or in gray color otherwise. RPM values were calculated as the average of two small RNA-seq replicates from for the same SH-SY5Y, MCF10A and MCF7 cells, and from an ENCODE HeLa cells. RPM values and qPCR values in ΔCt scale are given in Supplementary Tables S12 and S13. The qPCR experiment was evaluated by comparing each RNA expression with respect to the expression endogenous control U6 snRNA in each sample.

## Discussion

SeRPeNT provides a fast and accurate method to identify known and novel small non-coding RNAs exploiting read profiles from stranded size-selected RNA sequencing data. SeRPeNT does not depend on the annotation granularity and avoids many drawbacks inherent to sequence and secondary structure based methods, which may be affected by post-transcriptional modifications or limited by the reliability of structure determination. Here we have shown that read profiles, by capturing the post-transcriptional processing that is specific to each sncRNA family, provide functional information independently of sequence or structure. In particular, a number of known snoRNAs and tRNAs clustered with miRNAs according to their profiles. We also expect that our dynamic-time warping algorithm can account for the heterogeneity in the processing miRNAs (48) and other sncRNAs. Beyond the known cases, we detected new candidates of this dual behaviour. It remains to be determined whether these new sncRNAs can indeed function as miRNAs and associate with AGO2 (49). It is possible that they compete with more abundant miRNAs to be loaded on the RNA-induced silencing complex; hence they might become more prominent in specific cellular conditions. Incidentally, many sncRNAs increase expression when this is measured from the sequencing of AGO2-associated reads in *DICER1* knocked-down cells (data not shown), suggesting a repression by *DICER1* (33) or an association to alternative biogenesis pathways (39).

We have generated an extended annotation for human that includes hundreds of previously unannotated sncRNAs from known classes. These included new miRNAs, which we validated comparing to known families, confirming the structure of the precursor, and by measuring their expression dependence with the miRNA biogenesis machinery. We further observed the frequent differential processing of sncRNAs across cell compartments, especially for tRNAs. As differential processing of tRNAs has been observed in association to disease (50–52), the observed patterns may be indicative of relevant cellular processes that are worth investigating further.

We also detected 131 new sncRNAs that could not be labeled, which we named clustered uncharacterized RNAs (cuRNAs), and which are not present in current sncRNA catalogues, hence could correspond to novel sncRNA species. Although cuRNAs did not show frequent differential processing across cell compartments, they showed dependencies on the miRNA processing machinery and overlapped with CAGE tags or lncRNAs; suggesting mechanisms of biogenesis. The role of lncRNAs as possible general precursors of multiple types of sncRNAs in fact suggests new possible ways to classify lncRNAs beyond the current proposed frameworks (53). A subset of lncRNAs may act as precursors of a wide variety of small non-coding RNAs, including those from known families. On the other hand, cuRNAs conform a small fraction from all the known classes of sncRNAs, indicating that there might be a very limited number of new sncRNA species.

SeRPeNT is the first unsupervised tool for classifying and characterizing sncRNA processing profiles. Previous methods are supervised and need to rely on prior annotations in order to group and then classify novel sncRNAs. SeRPeNT does not need any annotation to cluster unknown sncRNAs and therefore is the only method capable of discovering novel sncRNA families and to group sncRNAs from newly sequenced organisms for which no phylogenetically close annotation exists.

We validated our approach by obtaining experimental evidence for the expression of four predicted sncRNAs, with no similarity to any other genomic locus, and from four different classes: one intronic miRNA, an snoRNA-derived miRNA, a processed tRNA and a cuRNA. Although we could experimentally validate the specific expression of these sncRNAs, we did not always find an agreement between the experimental validation and the detection by SeRPNT in the same cells. Some of the filters used for SeRPeNT might have been strict, thereby limiting our level of detection. Nonetheless, the validation of these new sncRNAs demonstrates SeRPeNT’s ability to detect RNA species that are experimentally reproducible. Further analyses and validations will be required to capture the extent and variability of the processing of these small RNAs across multiple conditions.

We envision a wide variety of future applications of SeRPeNT, including the fast identification and differential processing of non-coding RNAs from size-selected RNA-sequencing from tumor biopsies, circulating tumor cells, or exosomes, as well as the rapid discovery and characterization of non-coding RNAs families in multiple organisms. SeRPeNT differential processing operation can also be very powerful at, for instance, discovering RNAs that are differentially processed in tumor cells, thus generating biomarkers and potential drug targets. In summary, SeRPeNT provides a fast, easy to use and memory efficient software for the discovery and characterization of known and novel classes of small non-coding RNAs.

## Acknowledgements

We are thankful to D. Gautheret for useful discussions. EE and AP were supported by the MINECO and FEDER (BIO2014-52566-R), Consolider RNAREG (CSD2009-00080), by AGAUR (SGR2014-1121), by the European ITN Network RNP-Net (ID:289007) and by the Sandra Ibarra Foundation for Cancer (FSI2013). RG and AP were supported from MINECO (BIO2011-26205), National Human Genome Research Institute of the National Institutes of Health (U54HG007004) and MINECO Centro de Excelencia Severo Ochoa 2013-2017 (SEV-2012-0208).

